# Wireless recordings from dragonfly target detecting neurons during prey interception flight

**DOI:** 10.1101/2024.11.12.622977

**Authors:** Huai-Ti Lin, Igor Siwanowicz, Anthony Leonardo

## Abstract

Target interception is a complex sensorimotor behavior which requires fine tuning of the sensory system and its strategic coordination with the motor system. Despite various theories about how interception is achieved, its neural implementation remains unknown. We have previously shown that hunting dragonflies employ a balance of reactive and predictive control to intercept prey, using sophisticated model driven predictions to account for expected prey and self-motion. Here we explore the neural substrate of this interception system by investigating a well-known class of target-selective descending neurons (TSDNs). These cells have long been speculated to underlie interception steering but have never been studied in a behaving dragonfly. We combined detailed neuroanatomy, high-precision kinematics data and state-of-the-art neural telemetry to measure TSDN activity during flight. We found that TSDNs are exquisitely tuned to prey angular size and speed at ethological distances, and that they synapse directly onto neck and wing motoneurons in an unusual manner. However, we found that TSDNs were only weakly active during flight and are thus unlikely to provide the primary steering signal. Instead, they appear to drive the foveating head movements that stabilize prey on the eye before and likely throughout the interception flight. We suggest the TSDN population implements the reactive portion of the interception steering control system, coordinating head and wing movements to compensate for unexpected prey motion.

## INTRODUCTION

When confronted with moving target, a predator or pursuer can implement one of several different interception strategies. The simplest approach relies on reactive control, in which the pursuer responds to every change in the prey’s observed trajectory. This framework requires tremendous acceleration,^1,2^ fast reactions,^3–5^ and is lagged in time and thus error prone.^6,7^ A more sophisticated approach is to balance reaction and prediction, with predictive control handling the expected movements in the prey’s trajectory^8–10^ and reactions reserved for the unexpected. The cost of this model-driven strategy is an increase in sensorimotor complexity but with the benefit of less expended energy and higher success rates. Our prior work has established that dragonflies use this latter strategy, carefully balancing predictions of how the prey and its own body will move, with reactions to the prey’s sudden and unpredicted turns.^8^ Before flight begins, the dragonfly carefully observes the prey and decides if and when to pursue.^11^ When prey fly near a perched dragonfly, they trigger a rapid 50ms head saccade from the dragonfly that centers the prey in the dragonfly’s visual fovea, a high resolution region on the compound eye.^12^ Smooth pursuit head tracking then holds the prey image steady on the dragonfly’s fovea for ∼250 ms. Flight is triggered as the prey passes directly above the dragonfly. This interception lasts only a brief 300-600 ms. During the interception flight, the dragonfly continues to actively stabilize the image of the moving prey on its eye while keeping the prey directly overhead and progressively closing the distance (Figure 1A). As in many predatory animals,^4,6,10,13,14^ dragonfly prey capture is driven by vision – though precisely how remains unknown.

**Figure 1:**
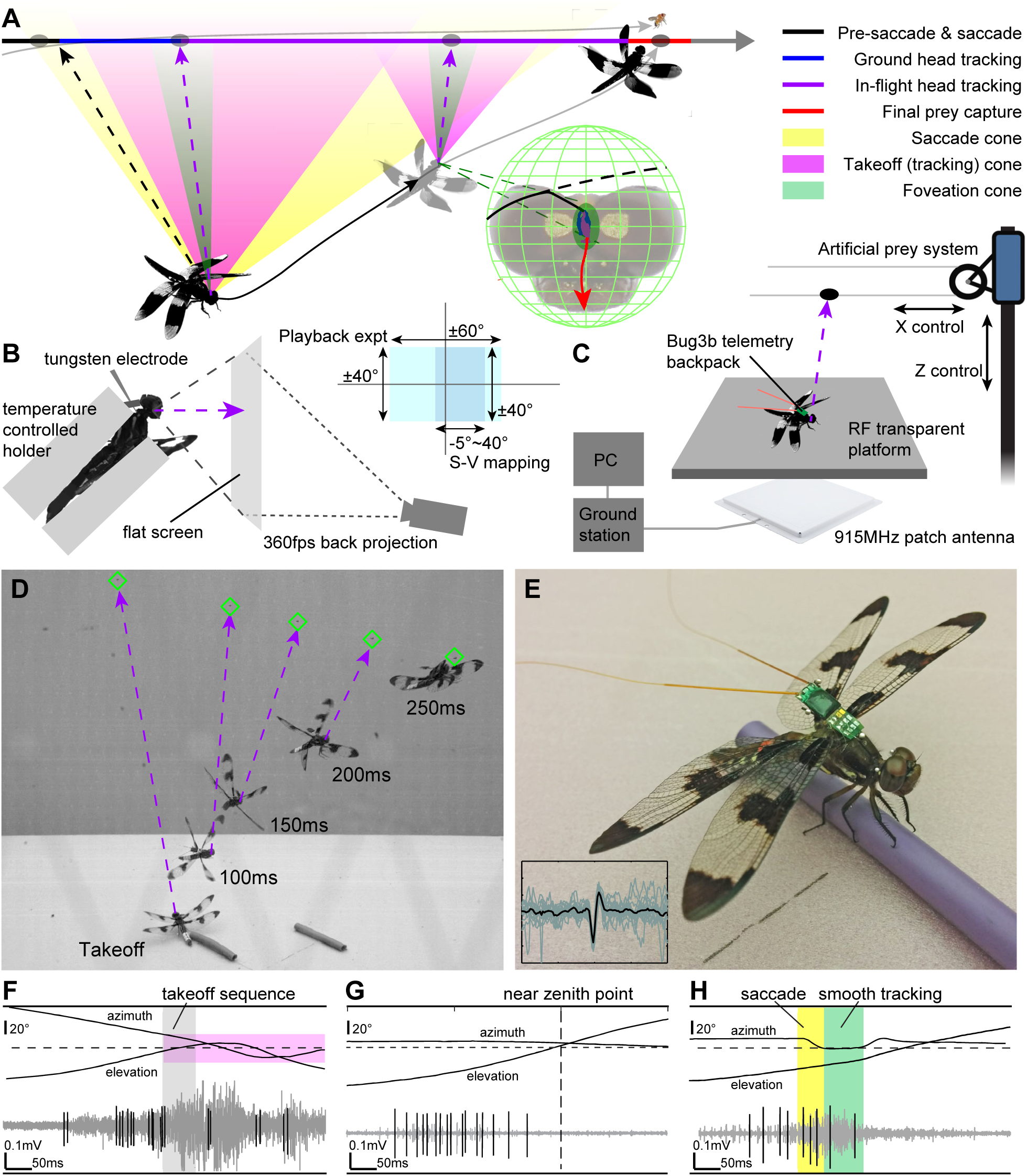
Dragonfly prey interception behavior and electrophysiological methods. **A)** Illustration of the dragonfly prey interception behavior (adapted from **Lin & Leonardo 2017**). The dragonfly makes an rapid head saccade to bring the target to the functional fovea, in which the target image wobbles until going downward into the mouth in the final capture stage. **B)** Electrophysiology recordings from immobilized dragonflies were supported by a temperature controlled dragonfly holder and a 360fps back projection. Two projection ranges were used for the target playback experiment (cyan) where real interception data were used and target size-speed mapping (dark blue) to accommodate different angular resolutions. **C)** Wireless electrophysiology experiments were achieved by combining the Bug3b telemetry (**Thomas *et al* 2012**) and an artifical prey system. **D)** An action sequence image of a dragonfly fully equipped with the neural telemetry backpack capturing an artificial target marked in green diamond. **E)** The dragonfly carries the backpack sampling at 26kHz or 52kHz. An example unit spike overlay is shown. **F)** Example prey interception telemetry recording with an isolated unit labelled, and the target angular position relative to the takeoff (tracking) cone. **G)** Example telemetry recording when the dragonfly made no movement to the artificial target given the corresponding target angular position relative to the head. **H)** Example telemetry recording when the dragonfly made a head saccade (yellow shade) and visually tracked the target (green shade) before giving up.

During prey capture, the dragonfly’s visual system encounters three different types of prey image motion. In the initial prey detection phase, the dragonfly is perched and its head is still; the prey image on the eye moves at roughly constant velocity along the flight path of the prey (straight lines with periodic turns). In the second phase of the behavior, the dragonfly remains perched but has foveated the prey via a head saccade^12^ and is performing smooth pursuit tracking of the target – the prey image drifts around the fovea at high speed with small displacements. In the final phase of in-flight pursuit, target foveation continues but prey image drift is driven not only by prey motion but also by the dragonfly’s movements. The dragonfly’s head motion actively cancels both disturbances via a combination of predictions and reactions.^8,11^ These different forms of prey image motion are significant for two reasons: first, to the extent that dragonfly visual neurons have been studied, they have been explored primarily with pre-takeoff constant velocity stimuli rather than those encountered in-flight. Second, and more significantly, if the prey image is largely stabilized on the eye during flight, it begs the question of what role visual neurons might play in interception – a profound question for a behavior that is fundamentally driven by vision.

All dragonfly species studied to date have a specialized class of 16 giant neurons that are hypothesized to measure the drift in the visual angle between the dragonfly and the prey.^15^ These target-selective descending neurons (TSDNs) individually are tuned exclusively to directional prey image motion on the eye.^15^ Collectively they tile the dorsal visual field, and overrepresent the fovea.^16^ Their population vector average has been shown to encode the direction of the target on the eye. This neural representation of prey velocity information has been noted in support of the notion that TSDNs implement a parallel navigation guidance strategy.^16,17^ It is hypothesized that TSDNs provide continuous steering signals, reactively derived from vision, to guide the dragonfly to the target much like a guided missile.^18^ Nonetheless, several key questions remain about the role of TSDNs in interception. First, while interception flight paths resemble parallel navigation, their fine-scale structure is inconsistent with it.^8^ Second, the connectivity of TSDNs to wing motoneurons (MNs), and therefore their ability to steer, remains unknown. Finally, TSDN studies to date have only been in immobilized dragonflies, watching constant velocity prey motion.^11,19^ TSDN responses during active head tracking and interception flight remain unknown. In this paper we use neuroanatomy and neural telemetry to characterize the role of TSDNs in prey interception and their contribution to the reactive and predictive control of steering.

## RESULTS

### TSDN responses to naturalistic prey motion

The TSDN recording preparation was pioneered by Olberg.^15,20^ A head-fixed immobilized dragonfly at room temperature (20-25°C) views visual stimuli on a screen while an electrode measures the spiking output of one or more neurons. TSDNs show no responses to wide field stimuli such as moving gratings. However, they are strongly driven by small moving targets (e.g. 1.5°∼3° in angular diameter).^15,16,20,21^ These targets elicit maximal responses when they move at constant speed in a cell-specific directionally tuned receptive field. This procedure has been the gold standard for functionally defining TSDNs. However, we now know that dragonflies rarely see constant velocity prey image movement during interception flights.^8^ We also know the angular size and speed of prey encountered during interception are considerably smaller and faster than those explored previously.^11^ It is thus difficult to hypothesize TSDN responses during behavior from those measured in immobilized animals. Such a comparison would be extremely useful, as it allows separation of the sensory and motor aspects of the response. We thus began our study by revisiting the immobilized recordings (Figure 1B) using ethologically correct stimuli in two experiments.

In our first experiment (n=5 dragonflies), we explored the effects of prey angular size (0.1°∼10°) and prey angular speed (100°/s - 1000°/s) on TSDN response strength. We covaried the dragonfly’s body temperature from torpor to flight temperatures (16°C, 20°C, 26°C, 32°C) to ensure we explored the full physiological regime; prior work has been conducted only at room temperature (i.e. 21°C). Extracellular TSDN responses were recorded while the immobilized, temperature-controlled dragonfly viewed a target moving at constant velocity. The angular size and speed of the target were changed randomly using the size and speed ranges described above (see Methods). We found the size-speed parameters that evoked high responses were a strong function of temperature. At low temperatures (16°C), TSDNs had weak responses and no responses to target speeds over 150°/s (Figure S2B,C). At room temperatures (20°C) the best stimulus was a target 1.5° in size moving at ∼200°/s speed (Figure S2B), consistent with traditional experiments.^20^ At high temperatures (32°C), TSDNs have strong responses to target size 0.5°∼1° and speeds up to the maximum we tested, 1000°/s (Figure 2A-B, 2D-E). The firing dynamics are fast, and we found short inter-spike intervals below 5 ms across all body temperatures (Figure 2C,F). Prior work has shown that foraging dragonflies take off after targets 0.15°∼0.7° in size traveling 100∼350°/s (Figure 2G). During the interception flight, these prey grow both in angular size and speed as the dragonfly approaches them (Figure 2H), but for the most part remain below 1° in size and below 350°/s in speed. When we plotted prey angular statistics at takeoff, we found they closely resembled the TSDN’s size-speed tuning profile (Figure 2I). In short, the firing rates of TSDNs at flight temperatures (32°C) match the size-speed statistics of natural prey encountered during target detection and interception flight.

**Figure 2:**
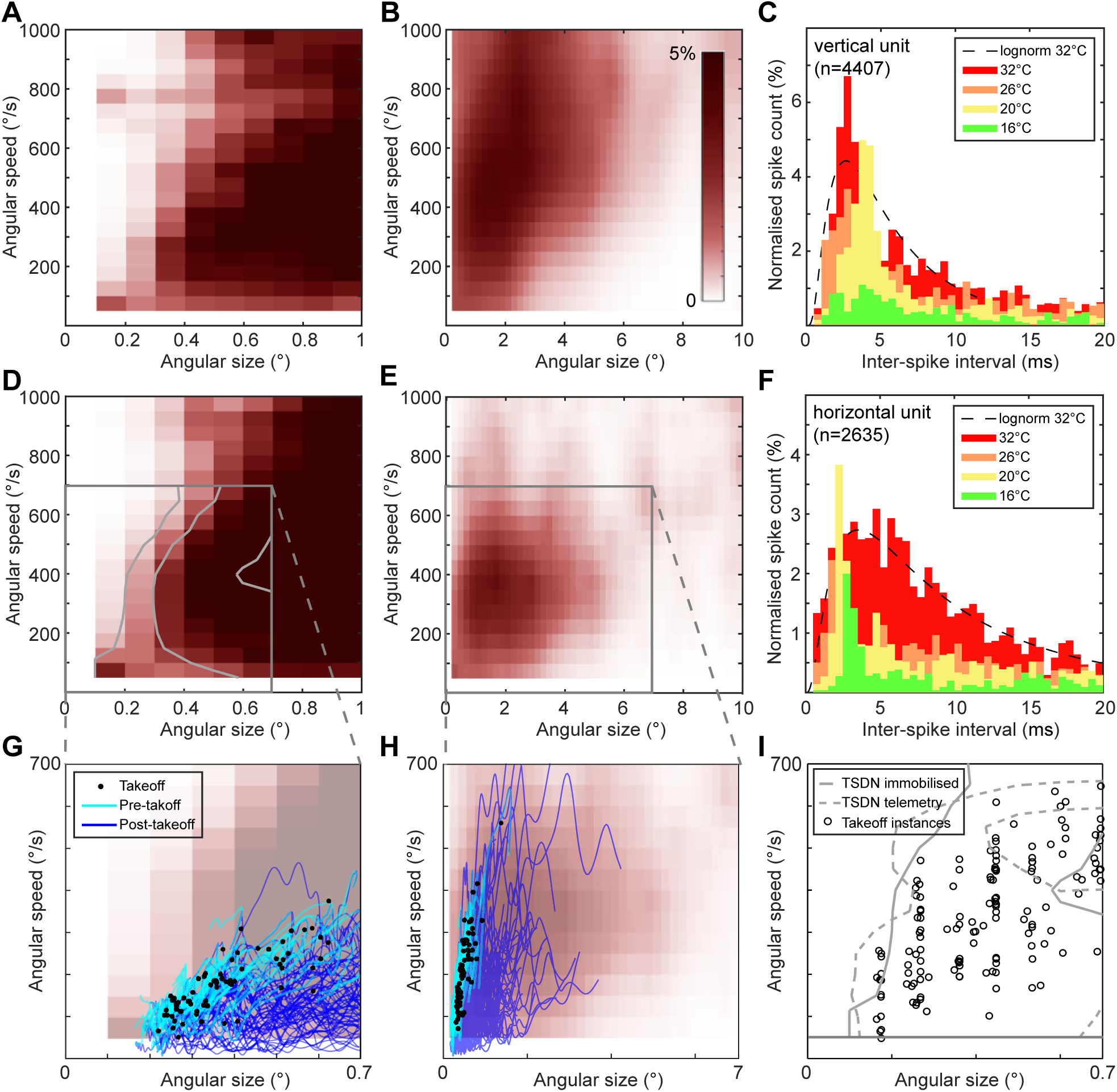
Target size-speed response tuning. **A-B)** The normalized spike count histogram of a TSDN unit responding to vertical upward moving target at 32°C thoracic temperature in two angular size mapping ranges. **C)** The inter-spike interval from mapping of the vertical unit at 16°C, 20°C, 26°C, and 32°C, normalized to the 32°C map spike count. All distributions show fast spiking events right against 1ms. Most spikes occurred within 5ms interval with peak around 3ms, capturing the bursty spiking nature. A lognormal fit was applied to the 32°C condition for reference (black dashed line). **D-E)** The size-speed tuning map of a TSDN unit responding to horizontally moving target at 32°C thoracic temperature in two angular size mapping ranges. **F)** The same plot as **C** for the horizontal unit shows similar peak response at ISI∼3ms. For all size-speed mapping, a single circular target was shown with a uniform distribution of target states presented at randomized order. The TSDN size-speed maps came from 6 immobilised dragonflies. **G-H)** The angular size-speed state evolution of real prey during the dragonfly prey interception events (n=75; data reprocessed from Mischiati *et al* 2014). Target trajectories are projected to the perch location (cyan), and to the dragonfly’s body position (blue). Black dots mark the time of takeoff. The size-speed maps from **D-E** are shown in the background for reference. **I)** Target size-speed distribution of pre-takeoff prey selection behaviour screening. The black circles mark the takeoff instance when the entire plot aera was uniformly tested for pursuit decision in our previous study (**Lin & Leonardo 2017**). Contours of TSDN spiking probability from immobilised mapping and telemetry recordings in this study show good agreement with the prey selection behavior.

In the second experiment (n=5 dragonflies), we explored how TSDN responses were influenced by the trajectory of the visual stimulus. In our previous studies, we measured both the prey and the dragonfly’s time varying position, as well as the dragonfly’s head orientation during interception flight. These data enabled us to create prey image that match the prey image trajectories seen during behavior. We refer to this as the “head-centered stimulus” to differentiate it from the conventional “perch-centered stimulus” in which the prey moves in a straight line with constant speeds. Perch-centered stimuli elicited a high frequency burst of spikes (Figure 3A,D) as the prey image entered the visual field. Head-centered stimuli, however, evoked a more complex response pattern (Figure 3B). The pre-foveation portion of the trajectory moved smoothly across large angular extents of the eye, much like a perch-centered stimulus, and elicited a strong TSDN response. Once foveation began, target image motion was restricted to the fovea, and had high angular velocity but small displacement. This latter type of target motion did not excite TSDNs effectively (Figure 3E). Consequently, TSDNs in the immobilized dragonfly responded primarily to the pre-takeoff component of the head-centered stimulus and minimally to the in-flight component.

**Figure 3:**
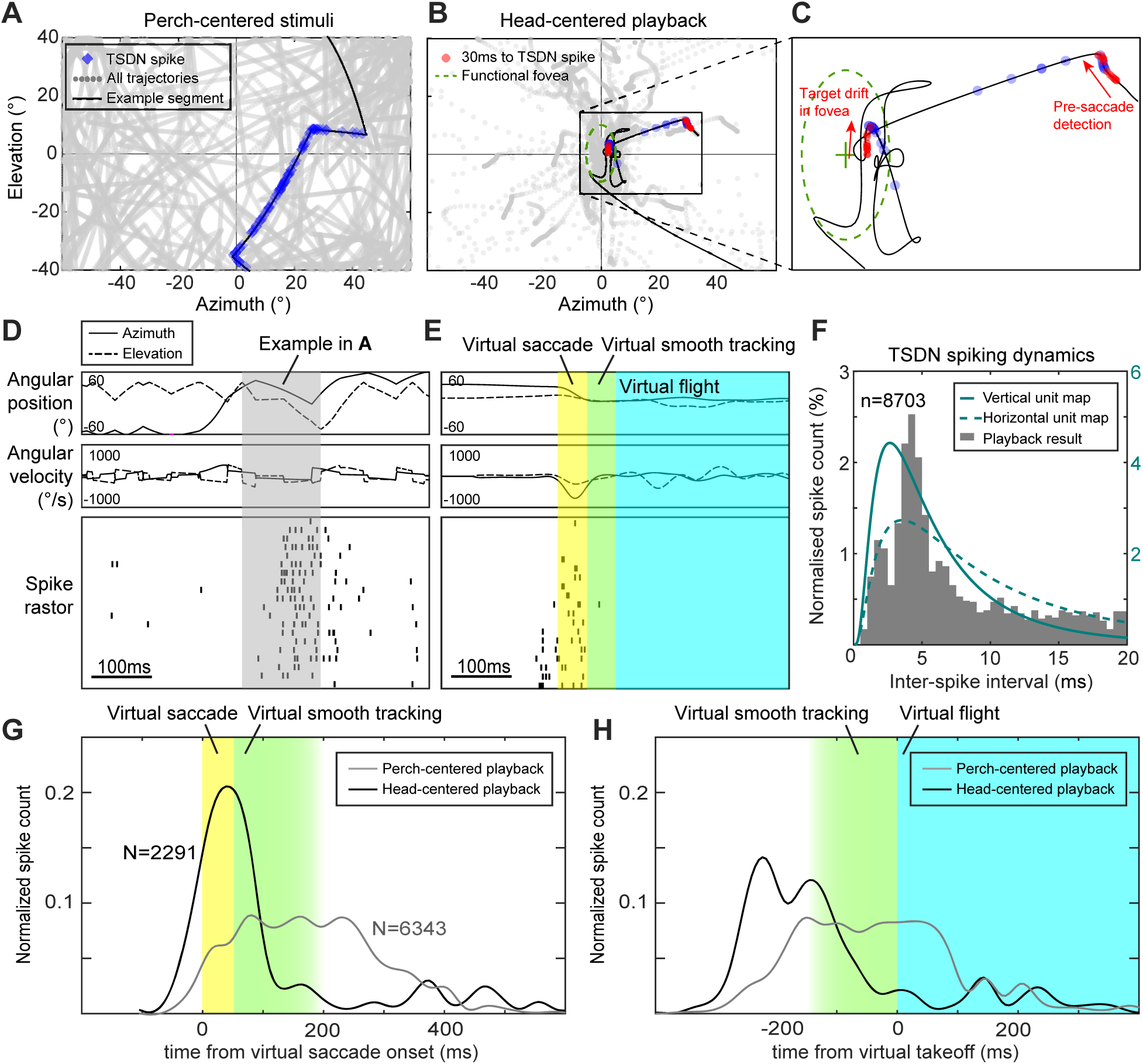
TSDN response timing to playback of prey trajectories. **A)** Classical linear constant veolcity target stimuli with random change of direction and speed projected to the angular coordinate. An example segment is highlighted in black with spike locations marked in blue dots. **B)** Real interception target trajectories playback projected to the head-centered reference frame which highlights the effect of active target foveation. The latency-corrected (30ms time shift) spike locations marked in red dots show that most spikes can be attributed to the pre-saccade trajectory, with some response to target drift in fovea. **C)** Zoomed-in image of the example in **B** shows the wobbly head-centered target trajecotry. Marking the spikes with 30ms latency in the past shows that TSDNs are responding to upward prey motion prior to the hed saccade and when targt drifts within the fovea. **D-E)** TSDN firing raster plot over 20 repeated stimulus presentations of one example trajectory from **A** and **B**. Cell adaptation is visible with decreasing response from early trials (bottom) to later trials (top). Yellow, green, and cyan shades represent the virtual head saccade, virtual smooth tracking, and virtual interception flight respectively. **F)** Inter-spike interval distribution for the entire playback experiment reveals the TSDN spiking dynamics. Overlaying the lognormal fits (olive green curves) from Figure 3F demonstrates the consistent fast spiking below 5ms. **G)** The aggregated spike timing histograms from head-centered playback (black line) or perch-centered playback (grey line). **H)** The same data in **G** aligned to the virtural takeoff. This dataset came from 5 immobilized dragonflies for the playback experiment.

The effect of foveation and takeoff can be summarized by the spike histogram aligned to the virtual saccade and virtual takeoff across all 5 dragonflies recorded (Figure 3G,H). It is well established^8^ that dragonflies maneuver throughout the interception flight. If these maneuvers are primarily driven by TSDN signals we would expect, on average, a relatively constant level of TSDN population activity during flight. Perch-centered TSDN response patterns are consistent with this hypothesis (Figure 3D). However, the visual stimuli encountered by behaving dragonflies is the head-centered stimulus, not the perch-centered stimulus. TSDN responses to head-centered stimuli suggest a very different role for these neurons. The head-centered TSDN spike histograms show a peak response during the target foveating head saccade (Figure 3G), and significantly more spikes prior to takeoff (Figure 3H). Once foveation is established TSDN activity was substantially reduced. In contrast, if we generated the stimulus assuming the dragonfly’s head was kept in the same orientation relative to the body, the target moved rapidly and often went outside of the receptive field soon after takeoff (Figure S3D). This “body-centered stimulus” produced a more sustained response in the TSDNs (Figure 3G,H). This implies that, given active head tracking, TSDNs will be largely quiescent during flight. The important caveat is that while the head-centered visual stimuli accurately represent those seen during interception, many other behavioral and sensory signals (such as proprioception) are absent in these immobilized preparations. It is thus crucial to have TSDN recordings in freely flying dragonflies hunting prey to validate these observations in immobilized animals.

### TSDN responses during hunting behavior

In-flight recording of TSDN responses have never been made for the simple reason that dragonflies, while large for insects, are exceedingly small to carry instrumentation. An adult *libellulid* dragonfly weighs 350-450 mg. In comparison, a miniature battery alone can weigh over 130 mg,^22^ and the mass of a full recording system can easily exceed the dragonfly’s own weight and lift capacity. To solve this problem, we engineered a micro-telemetry system suitable for recording neural signals from flying insects.^23^ A custom differential recording and ADC chip was developed to simultaneously measure up to 9 neural signals sampled at 26 kHz/channel. The chip was wire bonded to a small 9.5mm x 3.9mm flex board (Figure S1A). A miniature dipole antenna was developed to allow the recording system to backscatter the 5Mbps data stream and wirelessly harvest RF power to run the device (see Methods). The total mass of the telemetry backpack including motion capture markers was 45 mg, or <12% of a typical *Plathymis lydia* dragonfly’s body weight. We found that, carrying an inactive backpack alone, without electrodes, dragonflies could successfully forage and gain weigh over many days, indicating the payload and mounting configuration had no impact on successful predation.

Successful telemetry experiments were limited primarily by the electrode implantation surgery and secondarily by the precision backpack installation process (Figure S1B) with motion capture markers and electrode connections (see Methods). After excluding preparations with failed electrode implantation, roughly half the dragonflies were able to fly normally after the electrode-backpack implantation (14/30). 28% of these dragonflies engaged in prey interception flights (4/14). 75% of the prey intercepting dragonflies had sortable signals (3/4), and 25% of these hunting individuals captured the artificial targets successfully (1/4). The total yield of successful telemetry flights with usable in-flight TSDN data during prey interception was 11 flights from 3 dragonflies. More data were measured from telemetry dragonflies that perched and actively engaged in prey head tracking but did not take off (n=3 dragonflies, 138 events). Because of these challenges, we used neural telemetry to validate the results obtained in the immobilised preparations rather than as a primary data source.

Using the telemetry system, we first explored the role of TSDNs during the pre-flight head tracking behavior (Figure 1C,G-H). Consistent with the immobilized preparations, TSDNs were activated as the target entered and moved across the saccade cone (Figure 4A), with similar or even faster spiking dynamics (Figure S1C). We presented both real and artificial targets to perched, foraging dragonflies and monitored their pre-takeoff behavior -- a head saccade to foveate the prey followed smooth pursuit head tracking (Figure 4D, right). As noted earlier (Figure 3), during the pre-saccade regime the prey image on the eye is approximately a perch-centered stimulus, similar to those used in immobilized experiments, while after the saccade active head tracking hold the prey image steady on the eye. During these perched head tracking events, the telemetry signal generally contained 1-2 small target related units (Figure 1G,H). While we used standard criteria to define these units (see Methods), it was often difficult to reliably sort them into identified cells (e.g. MDT1). We structured our analyses so none of the results to follow are dependent on whether the unit was perfectly isolated or partially multi-unit. When a foveating head movement occurred, the initial burst of spikes was interrupted and rapidly quenched by the head saccade (Figure 1F,H; Figure 4D, right). TSDN firing events then remained rare throughout the foveation period. The TSDN spike response histograms indicate that the probability of firing increased quickly when the target appeared, ∼40ms prior to the head saccade, and peaked around the head movement (Figure 4F, n=3 dragonflies, 138 events). Firing rates then decreased monotonically until 200ms post head saccade onset. This is consistent with the TSDN response to playback of head-centered prey trajectories with virtual head saccades (Figure 3G) and is the first quantitative link between TSDN responses in the immobilized and behaving dragonfly.

**Figure 4:**
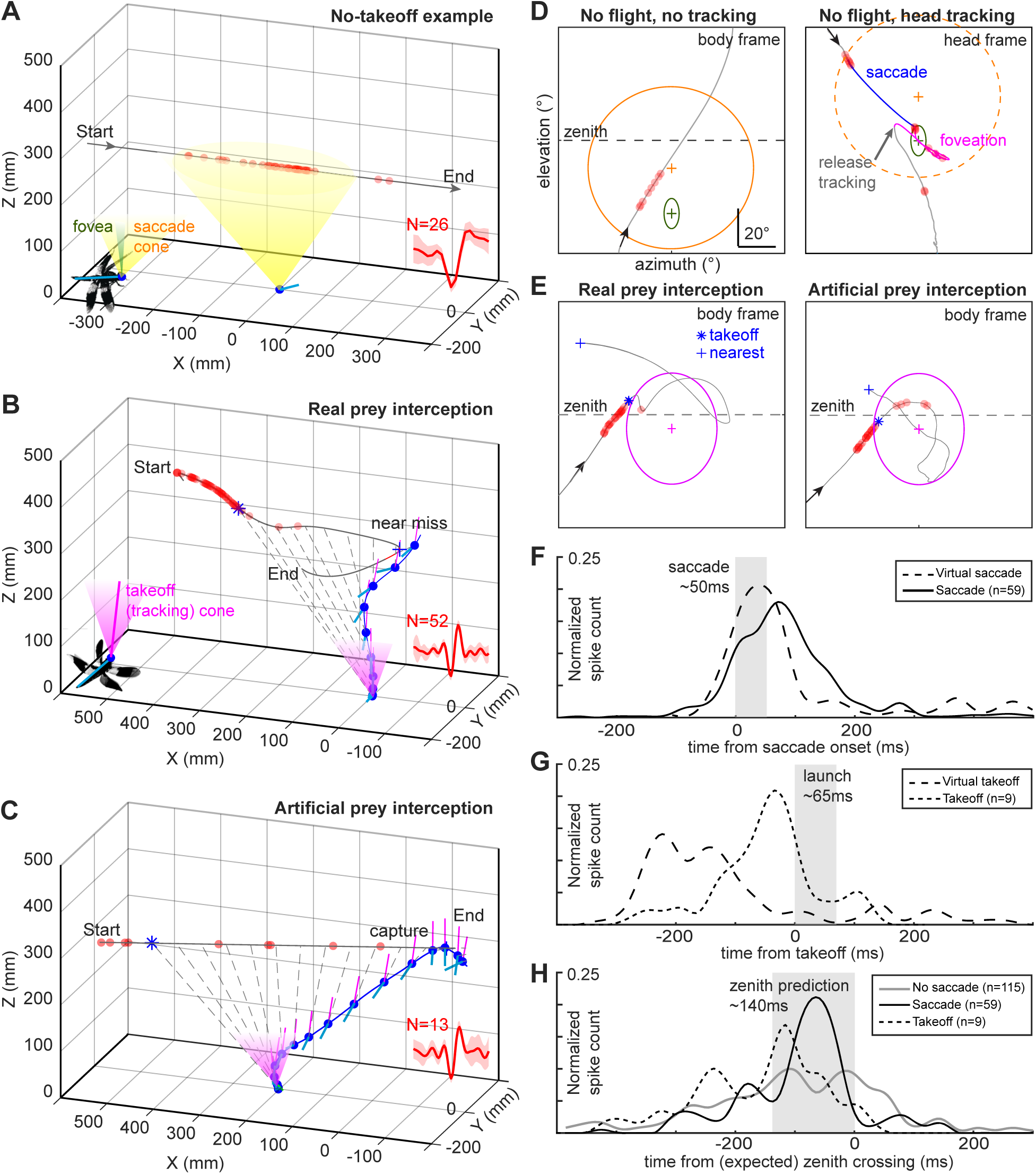
Target responses from hunting dragonflies. **A)** An example of an artificial target going through a hunting dragonfly’s visual field without a pursuit flight. The target responses (red dots; waveform overlay as inset) started promptly as the target entered the saccade cone (yellow) of the perched dragonfly (blue dot and cyan stick as illustrated). **B-C)** Example interception of real prey and artificical prey with takeoff (blue *), and nearest point (blue +) marked. The line-of-sight vectors are shown in 50ms intervals (dashed grey lines). The dragonfly rotated its body to align to the target direction immediately after takeoff, while trying to keep the target within the tracking cone (magenta, as illustrated). The waveform overlay is shown as insets. **D)** Examples of the target responses when the dragonfly did not take off without the foveating head tracking **(left)** and with head tracking **(right)**. The saccade cone (orange ellipse; 75° elevation) and functional fovea (green ellipse; 52° elevation) are marked. The saccade cone is reference to the body thus becomes irrelavant once head tracking started. **E)** Examples of target responses of dragonfly intercepting a fruit fly (left) and an artificial prey (right). The takeoff events were consistently triggered (blue *) as the target entered the takeoff (tracking) cone (magenta ellipse). **F-H)** Spike timing histogram aligned to head saccade, takeoff, and zenith crossing. Conditions are labelled in the legends.

We also used the telemetry system to measure the TSDN prey angular size-speed tuning while perched dragonflies observed flying prey (3 dragonflies, 174 events, 514 spikes). This was done opportunistically when the dragonfly’s head was in a relatively reproducible orientation (±30°), by accumulating TSDN responses to artificial prey prior to head saccades. This target size-speed map measured with telemetry (Figure 2I) shows a tuning contour consistent with that seen in the more detailed immobilized experiments (Figure 2D) as well as with the behaviour take-off tuning previously reported.^11^ In this previous study, we uniformly presented every combination of target angular speed and size (within predefined ranges) to more than 9 freely hunting dragonflies without any manipulation. The target states that were selected for interception flight (Figure 2I, black circles) closely match to the TSDN responses maps from both the immobilized dragonflies and telemetered dragonflies prior to head saccades.

Finally, we asked how TSDNs respond during interception flights (n=3 dragonflies, 11 flights; Figure 4). All interceptions of real prey (n=8 flights) by telemetered dragonflies ended unsuccessfully in the final moments of the flight when the dragonfly attempted to grab the prey in the air with a coordinated movement of its legs (Figure 4B; see Methods for a discussion). Several interceptions of the artificial targets (n=3 flights) were successful (Figure 4C). Consistent with the head-centered stimulus datasets (Figure 3), TSDN responses during full interception flights started well before takeoff as the prey entered the dragonfly’s field-of-view (Figure 4G). Takeoff generally occurred as the prey approached the zenith,^11^ and entered the takeoff (tracking) cone (Figure 4E). The maximum TSDN response during these interception flights occurred ∼50ms before takeoff, and then rapidly grew quiet for the remainder of the flight. There was no indication that TSDNs continuously steer the dragonfly nor continuously report the prey’s line-of-sight relative to the dragonfly. Collectively, TSDN recordings from hunting dragonflies support our findings from the immobilized preparations. The data are consistent with the idea that TSDNs detect prey, drive foveation, and likely report occasional degradation of target tracking before or during prey interception flight.

### TSDN connections to wing and neck motoneurons

If TSDNs drive portions of the interception flight, they require connectivity to the motor system. We asked whether TSDNs directly synapse onto wing or neck motoneurons (MNs). Such connections have been speculated for decades,^24^ but have never been confirmed. We backfilled TSDNs in bulk through a cut neck connective using 3kDa dextran conjugated with Texas Red. Because fluorescent dextran’s diffusion is significantly slower in smaller diameter processes, labelling was heavily biased towards the uniquely large 20-25um diameter TSDN axons. In the same specimen we performed multiple retrograde fills through motor nerves dissected from individual flight muscles of both the fore- and hindwing, using Neurobiotin as a tracer (Figure 5A-B). By dissecting the nerve branches from the muscles, we could identify the flight MNs without ambiguity.

**Figure 5:**
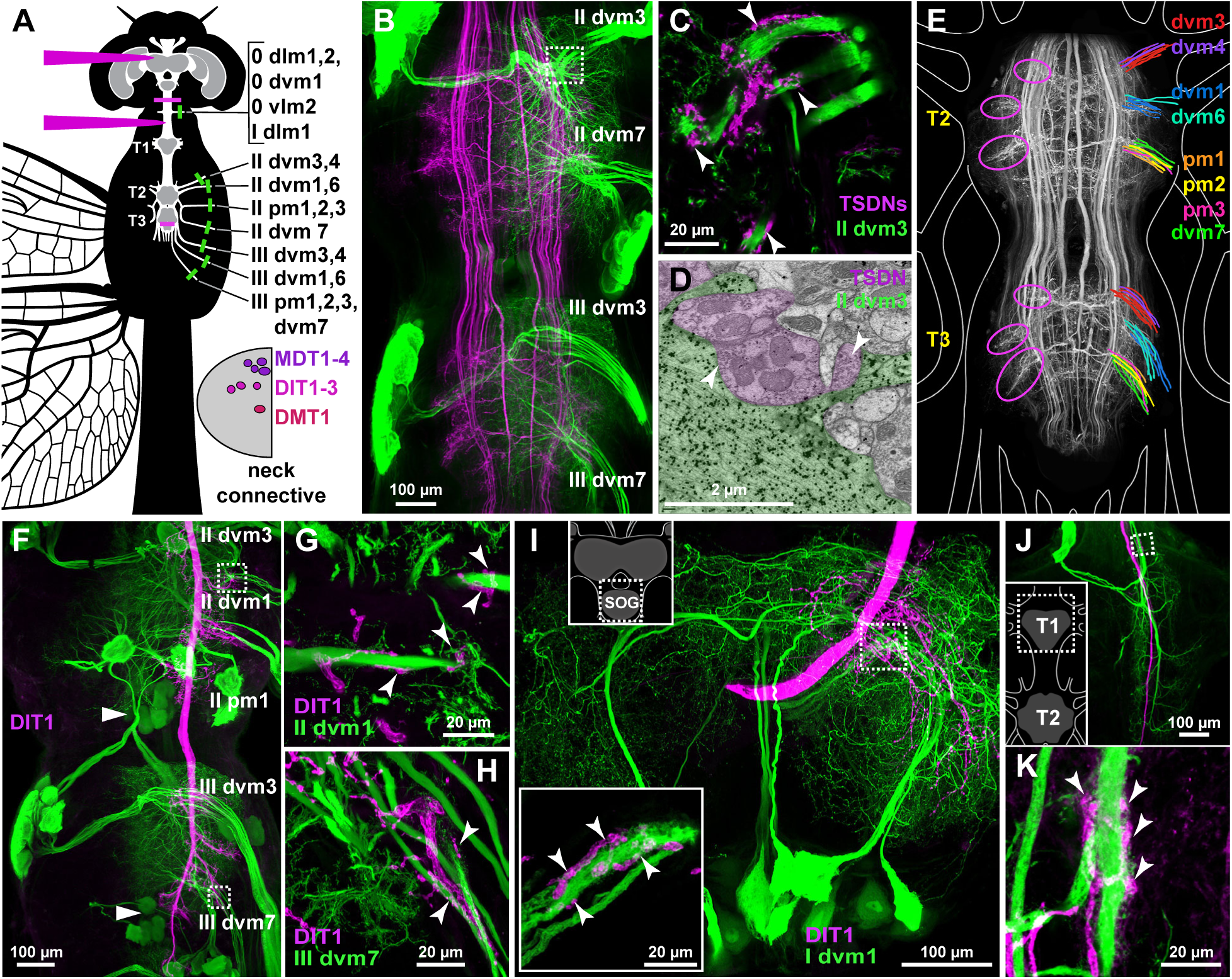
TSDNs synapse directly and extensively onto wing and neck muscles MNs. **A)** A diagram showing the dragonfly CNS (T1 – prothoracic ganglion; T2 – mesothoracic; T3 – metathoracic) and delineating experimental approach to TSDNs and MNs labeling (Methods). Color represents the tracer used to perform the fills: magenta – Texas Red-dextran; green – neurobiotin. Bottom right: location of TSDNs axons within the neck hemi-connective (Methods). **B)** Maximum intensity projections show the bulk-backfills of TSDNs (magenta) and MNs innervating wing muscles (green). **C)** Zoomed-in of white square in (B) showing contacts of TSDNs axonal terminals and wing muscles MNs trunks. Arrowheads point to several putative contact sites. **D)** Electron photomicrograph demonstrating direct connectivity between a TSDN (purple – 6 nm gold nanoparticles) and the wing muscle MN II dvm3 (green – 12 nm). Note the presence of vesicles in the TSDN’s profiles (white arrows). **E)** A diagram of TSDNs axonal output domains (magenta ovals) and wing muscles MNs tracts as they exit the T2 and T3 neuropiles. **F)** Single TSDN DIT1 labeled (magenta) and several wing muscles MNs (green). Arrowheads point to octopamine-immunoreactive cell bodies of unpaired median neurons. **G, H)** Zoomed-in of the white squares in (F). **I)** Suboesophageal ganglion with labeled DIT1 and MNs innervating right head elevator muscle I dvm1. Contacts are highlighted in inset. **J)** Prothoracic ganglion (T1) from the same preparation with two I dvm1 MNs in green. **K)** Projections showing the MN’s axon wrapped by DIT1 axonal terminals.

Confocal microscopy of the bulk fill preparation revealed a striking pattern (Figure 5E): the axonal terminals of TSDNs are restricted to six distinct domains of dorso-lateral meso- and metathoracic neuropil (3 per side, per ganglion). These output domains occupy the same space as and are adjacent to the dendritic trunks and axons of the wing MNs: The anterior-most domain contains MNs for dvm3 and dvm4; the middle domain contains MNs for dvm1 and dvm6; the posterior domain contains dvm7 and pm1, 2 and 3. Furthermore, the heavily beaded TSDN axonal terminals wrap around wing muscle MN trunks and follow them to the neuropil’s boundary (Figure 5C). The trunks are densely covered in dendritic processes that intermesh with TSDNs terminals (Figure 5C), forming the only points of contact between TSDNs and wing MNs. In contrast, the large and dense dendritic arbores of the wing MNs do not form any connections with TSDNs. This innervation pattern is unique among insect nervous systems. To verify direct connectivity, we used antibodies (against Texas Red and Neurobiotin) labelled gold nanoparticles of 2 different sizes, which were microinjected into two TSDNs (MDT, DIT), and MNs of dvm3 and dvm7. Immuno-electron microscopy validated contacts between the profiles of TSDNs and MNs terminals, as highlighted by 6 and 15 µm gold nanoparticles respectively (Figure 5D). We observed large quantities of vesicles in the profiles corresponding to TSDNs, and several densities at the sites of the two profiles’ contacts that could be interpreted as synapses (although morphologically different from the characteristic dipteran T-bars). Based on this EM evidence, we felt confident in using subsequent confocal data to identify TSDNs-MNs connectivity.

To further study the relationship between TSDNs, wing MNs and neck MNs, we next pressure-injected the dextran tracer into individual TSDNs either via the optic glomerulus or the neck connective (Figure 5A) while concurrently labelling several sets of wing and neck MNs (Figure 5F-K). We chose wing muscle MNs combinations that use non-overlapping tracks (Figure 5E), hence avoiding ambiguity in identifying target MNs. The identity of TSDNs was determined from the confocal data using approach described in Methods. The musculoskeletal organization of dragonfly’s neck and head was previously described.^25–27^ Forward-fill through a cut cervical nerve labels axonal terminals of MNs controlling four muscles: 0 vlm2, 0 dlm1, I dvm1 and I dlm1. These four muscles are controlled by MNs originating in both SOG and prothoracic ganglion (T1).

From this pressure-injected preparation, we found that **1)** the TSDN-MN connectivity pattern is indistinguishable between T2 and T3; **2)** bilateral TSDNs make contacts with both left and right MNs to similar degree; **3)** unilateral TSDNs sensitive to lateral translation of the prey contact MNs of dvm7, dvm6, and dvm3; **4)** the main power muscles, dvm1 and pm1, receive the least amount of the TSDN input; **5)** TSDNs make extensive and large-scale contacts with trunks of the MNs that supply neck muscles via cervical nerve. Their axons wrap tightly around MN trunks as they are about to exit the SOG neuropil (Figure 5I) or T1 neuropil (Figure 5J-K), and follow them into the neck connective. **6)** Double bulk-backfill of TSDNs and oblique neck muscles MNs (0 ism1-4) showed lack of evidence for direct connection. In conclusion, many individual TSDNs directly innervate both the neck and wing motoneurons in the dragonfly.

## DISCUSSION

Our study reshapes our understanding of the role of dragonfly TSDNs during interception. TSDNs were discovered 40 years ago.^15^ They were immediately recognized as a crucial neural locus for interception. In the years since, a steady body of work has focused on them as the primary means of interception steering.^16,20,21^ However, these hypotheses were grounded exclusively in data from immobilized animals. Here we have recorded from TSDNs during interception flights for the first time (Figure 1D-H, Figure 4) and clarify with several lines of evidence the role of these neurons in prey capture. We showed that the tuning of TSDNs to prey angular size and speed in both immobilized and behaving dragonflies (Figure 2) is well matched to the prey selection statistics seen in the behavior study of the same species^11^ (Figure 1A). We also show, for the first time, the anatomical wiring of TSDNs to both neck and wing motoneurons (Figure 5). Finally, we show that while TSDNs respond strongly to perch-centered constant velocity prey moving seen prior to takeoff, they are only weakly responsive to the foveated prey seen by the dragonfly during flight (Figure 3). Together these observations strongly argue against TSDNs as the primary steering substrate in interception, and instead suggest they play a core role in head steering, prey foveation, and the coordinated response of the head and body to unexpected prey movement. We discuss the implications of these points below.

The original hypothesis for TSDNs in interception has been inspired by guidance laws like parallel^28^ and proportional navigation.^3,4,29,30^ In these formalisms, changes in the line-of-sight angle of the target relative to the pursuer are used to update the pursuer’s interception steering trajectory. TSDNs were believed to provide the prey angular position and velocity needed to implement such guidance. Our data paint a more complex picture. We have shown that TSDNs do not simply drive the wings but also play an important role in coordinating the wings with the head. Such tight head-wing coordination is absent from the model of TSDNs as continuous steering units for interception flight.^16^ More significantly, prey foveation quenches prey motion on the eye, and the roughly constant velocity paths seen prior to takeoff are transformed into low amplitude high frequency jitter centered on the fovea. We have shown that TSDNs respond very little to these foveated prey images, both in immobilized and behaving dragonflies (Figure 3,4); only a sudden turn by the prey would elicit sufficient visual drift of the prey image and a TSDN response (Figure 3B-C, Figure 4D-E). The hypothesis that TSDNs serve as a flight autopilot based on a population vector average^16^ thus appears incorrect; there is not nearly enough visual stimulation or TSDN activity to support a continuous steering signal. One might ask if a separate feedback control system is used to maintain the “steady” interception trajectory, with TSDNs stepping in only when unexpected prey motion merits a correction. However, we have already shown this is not the case.^8^ Even when the prey is not maneuvering, the dragonfly’s flight path is not on a fixed heading and the dragonfly is continually maneuvering as a result of a series of internal model predictions about expected prey motion.^8^ These latter actions encompass the bulk of steering and appear to be entirely unrelated to TSDN activity. TSDNs must therefore play a different role than that of the principal steering channel.

The remarkable anatomy that wires individual TSDNs to both neck and wing muscles suggests TSDNs play a key role not in steering but in prey foveation. The single best stimulus to elicit a large TSDN response is prey moving at constant speed, with an angular size and speed closely matched to what a dragonfly observes shortly before takeoff. Such tuning is largely consistent with the contrast sensitivity theory for target detection.^31–33^ The extensive synaptic connections between TSDNs and the neck motoneurons would in principle allow them to directly rotate the head to stabilize the prey image on the eye. Indeed, we observed peak TSDN firing rates well before takeoff, in the period of constant velocity prey motion immediately prior to the foveating head saccade (Figure 3,4). In this non-flight period, the synapses of TSDNs onto wing motoneurons would presumably be ineffectual as the flight motor is off. However, once in flight a sudden movement by the prey, out of the fovea, would drive a TSDN response that could simultaneously adjust the state of both the head and the wings. We hypothesize this action re-foveates the prey and makes corrections to the flight path. Notably, those corrections would be brief and compact in time, as re-foveation quickly eliminates the visual signal driving the TSDNs; but it would be a small nudge in the corrective direction. Thus, we conclude that TSDNs are functionally (Figure 2,3,4) and anatomically (Figure 5) well positioned to drive the rare reactive responses needed to handle unexpected prey motion and maintain foveation. They may contribute to planning the flight path before takeoff, but the bulk of real-time interception steering is likely not driven by vision.

A core principle in modern sensorimotor control is the use of predictive internal models to plan and execute movements.^34–37^ Forward models predict the sensory consequence of a movement, and inverse models predict the motor commands needed to achieve a desired sensory outcome. In our previous work,^8^ we showed behaviorally that dragonflies rely on both forward and inverse models to generate interception trajectories and speculated that such models can operate in the absence of vision using internal state variables to update the flight path. The neural implementation of these models remains unknown, and an important question is whether they are generated by circuit dynamics, learned patterns of synaptic connectivity or both. While many studies clearly implicate circuit dynamics as a key element of predictive models in mammals,^34,35,38^ our study has led us to speculate on the role synaptic connectivity might play in directly implementing the inverse model as a lookup table. Reactions to foveal prey-image drift are relayed by TSDNs to neck and wing motoneurons and must be transformed into the motor commands needed to re-foveate the prey^8^ (Figure 5). The TSDN anatomy we have discovered is highly specialized. Instead of being integrated by the neuropil with other interneurons as in other systems,^39–41^ TSDN synapses occur on the trunks of motoneuron dendrites. This type of connection is suggestive of a fast modulatory function and has not been reported in any other system to the best of our knowledge. In short, TSDN signals are intrinsically sensory signals that must be converted to a motor command, but their MN synapses after the neck and flight motor circuit dynamics (e.g. flight CPG). Based on this, we speculate that the inverse model could be directly embedded in the TSDN-neck and TSDN-wing motoneuron synaptic weights, with these weights largely learned via evolution and then fine-tuned in the behaving animals. Exploring the plausibility of synaptic inverse model in the dragonfly is a topic for future studies.

Our findings resonate with recent advances in insect sensorimotor control. Hoverfly TSDNs, believed to detect and track conspecifics,^42^ respond sparsely to playback of simulated conspecific trajectories.^43^ Similarly, a brief single spike from the giant fibre^44^ (found in many dipteran species) directly drives leg motor neurons to trigger a low-latency escape jump independent of the normal takeoff control module.^45^ Giant fibres and TSDNs appear highly analogous in both anatomy and function – both fire sparsely and induce a brief and rapid modulation of an ongoing motor state rather than steer or navigate the animal continuously. In contrast, continuous steering neurons have been found separately. Many breakthroughs in insect descending neurons have emerged in the last decade via newer methods such as connectomics and optogenetic screening.^46–48^ A population of descending neurons in Drosophila, DNg02, directly regulate the wing stroke amplitude during flight, controlling the steering and thrust.^49^ Activity in these cells is not sparse and closely tracks wing beat amplitude stroke by stroke. This separation of fast reactive pathways and slower continuous control pathways may be a universal architectural feature of sensorimotor control. To keep the sensory system peeled for external stimuli and to enable model-based predictive control, efference copy could play an important role.^35,38,50–52^ The extremely fast, sparse, and bursty dynamics of TSDNs are reminiscent of ‘sparse coding’ observed in the putative upstream partners Small Target Motion Detectors (STMDs) and in vertebrate nervous systems.^53–56^

In a series of three papers, we have systematically revealed how dragonfly prey interception is implemented, through guidance algorithms, neuroanatomical evidence, and electrophysiological measurements. Our first study^8^ showed dragonflies employ a balance of reactive and predictive control. Forward models convert an efference copy of the flight motor plan into a body state prediction. An inverse model then integrates this prediction with the measured and predicted prey motion to adjust the prey foveating head command. Our second study^11^ dissected the foveation dynamics and showed that most critical prey parameters are carefully assessed prior to takeoff and used to plan the flight. Foveation quality is the most important of these parameters and dictates the prey pursuit decision as well as the interception success. In this third study, we measured the well-known target sensitive descending neuron activities directly in freely hunting dragonflies and demonstrated that they are carefully tuned for naturalistic prey motion and are most active prior to takeoff rather than in-flight. TSDNs are thus well suited to drive the head foveation and can simultaneously modulate both the head and flight motor system via direct motoneuron innervation. Based on these results, we speculate that many of the internal models used by the dragonfly are implemented via the synaptic weights between TSDNs and motoneurons of the neck and wings. Collectively, our three studies have progressed our understanding of aerial prey interception beyond the phenomenological level, producing a mechanistic framework for future investigations.

## Acknowledgements

We would like to thank Wade Sun for preparing custom electrodes and assembling telemetry backpacks. We thank Janelia Instrumentation Design and Fabrication department between 2012 and 2016 for providing generic engineering support for the project. We thank David Parks and the Janelia vivarium for dragonfly husbandry. We thank Prof Holger Krapp and Prof Gaby Maimon for their constructive criticisms to the manuscript. We thank Dr William Mowrey and Dr Matteo Mischiati for their discussions over many lab meetings. We thank Dr Daniel Ko for implementing the template matching algorithm into a GUI: DragonSort. The experimental work was supported by the Howard Hughes Medical Institute. Data analysis was supported by European Research Council (ERC-StG no.804315 ‘Vision-In-Flight’ to HTL).

## Author contributions

H-T.L. and A.L. designed and developed the acute and wireless electrophysiology experiments. H-T.L. performed the behavior and electrophysiology experiments. H-T.L. analyzed behavior and electrophysiology data. I.S. and A.L. developed neuroanatomy experiments. I.S. performed tissue preparation, imaging and analysis of the anatomical data. H-T.L. and A.L. wrote the original manuscript with input from I.S. All authors reviewed and edited the final manuscript.

## MATERIALS & METHODS

### Insect husbandry

A single dragonfly species *Plathemis lydia* was used for this study. Mature adults of both sexes were used in this study as we found no differences in prey interception behavior between the sexes. The insect husbandry procedures can be found in our previous publications.^1,2^

### Labelling of TSDNs and motoneurons

Dragonflies were anesthetized on ice, fixed on Sylgard-filled Petri dishes (Sylgard silicone elastomer kit, Dow Corning) using fine pins. For labelling of wing and/or neck muscles motoneurons through the cut ends of their axons, the insects were mounted right side up and the right ptero- and/or prothoracic pleural wall was removed. Motoneuron axon bundles were isolated/exposed by peeling away muscle fibres, placed in a drop (∼0.7 µl) of distilled water inside a petroleum jelly well, and cut. Water was wicked away, replaced with ∼0.5 µl of tracer solution (2 % neurobiotin, Vector Labs) and the wells were sealed closed with petroleum jelly. To simultaneously label TSDNs, right half of the prothoracic cuticle was removed, exposing neck connective, which was then cut and placed in a drop of water inside a petroleum jelly well. Water was quickly replaced with 5% 3kDa Texas red dextran (Life Technologies) in dH2O. When labelling neck muscles MNs, an alternative route for bulk-filling of TSDNs was used; in those cases, the T3-A1 (metathoracic-first abdominal ganglion) connective was cut through instead. Throughout the procedure, the tissues were sparingly washed with dragonfly saline (134 mM NaCl [7.83 g/L], 5.4 mM KCl [0.4 g/L], 3.8 mM CaCl2 [0.42 g/L], 3 mM MgCl_2_ × 6H_2_O [0.61 g/L], 25 mM Sucrose [8.5 g/L]…) to prevent them from drying and facilitate visibility, essential for spotting the translucent MN axons. Animals were kept refrigerated in a humid chamber for ∼20 hours to allow diffusion of the dye.

Labelling of single descending interneurons was accomplished by high-pressure microinjections into their axons in the neck connective or into their dendrites in the optic glomerulus. Standard-walled borosilicate glass capillaries were pulled and broken back by rubbing against or pushing through Kimtech lint-free tissue paper to increase the diameter of the opening, resulting in the ID at the tip of 10-15 μm. The capillaries were back-filled with 0.5 μl of 5% Texas Red dextran, coupled to a pressure pump (WPI model PV 820) and the pump was calibrated by injecting a droplet of the tracer solution into mineral oil; the pressure and duration were adjusted to allow the extrusion of ∼1 nl (corresponding to a spherical droplet of ∼125 μm). The central nervous system of the dragonfly is wrapped in tough and robust glial sheath that must be locally removed prior to injection to allow the micropipette’s entry. We used a high gauge hypodermic needle bent at 90° angle at the very tip to form a tiny hook, with which we lacerated and peeled off a small fragment of the sheath at the injection site. Typically, 1-2 nl were injected and the tracer was allowed to diffuse for 12 hr for VNC and 24 hrs for optical glomerulus at 4°C.

### Sample preparation

We adapted the fixation, permeabilization and clearing technique/protocols from Ott (2008), with some modifications. After dye diffusion ganglia were removed under HEPES-buffered dragonfly saline (HBDS; formula modified from Ott to match osmolarity of dragonfly saline: 134 mM NaCl [7.83 g/L], 5.4 mM KCl [0.4 g/L], 3.8 mM CaCl2 [0.42 g/L], 3 mM MgCl_2_ × 6H_2_O [0.61 g/L], 25 mM Sucrose [8.5 g/L], 10 mM HEPES [2.38 g/L], 0.03% Triton-X100) and subsequently fixed in 2% paraformaldehyde in Zn-HBDS (110 mM NaCl [6.5 g/L], 5.4 mM KCl, 1.5 mM CaCl_2_ [0.165 g/L], 3 mM MgCl_2_, 18.4 mM ZnCl_2_ [2.5 g/L], 0.5 mM NaHCO_3_ [0.04 g/L], 10 mM HEPES, 25 mM Sucrose, 0.03% Triton-X100) for ∼12 h at 4°C. Ganglia were then washed (2 × 20 minutes in HBDS, 2 × 20 minutes in PBS-T [phosphate buffered saline, 0.03% Triton-X100]) and, in order to make the glial sheath permeable, treated with a cocktail of proteases (0.25 mg/mL collagenase/dispase [Roche #10269638001], 0.1 mg/mL proteinase K [Roche #03115879001] and 0.25 mg/mL hyaluronidase [Sigma Aldrich #H3884-100MG] in PBS-T) for 30 minutes at 37°C. The samples where then washed with 100 mM Tris HCl buffer, 2 × methanol and finally incubated for 2 h at room temperature in 80% methanol, 20% DMSO to further permeabilize the tissues. Following that, the tissues were re-hydrated in 100 mM Tris and stained with DyLight 488-NeutrAvidin (1:250, Thermo Scientific #22832) in PBS with 3% normal goat serum, 1% triton X-100, 0.5% DMSO and Escin (a surfactant from the saponins family; 0.05 mg/ml, Sigma-Aldrich #E1378) at room temperature with agitation for 2 days. Long incubation times and the presence of surfactants assured better penetration of NeutrAvidin into the relatively thick and compact tissue of the ganglia. The samples were then washed in PBS/1% Triton and post-fixed for 4 hours in 2% PFA at room temperature. To avoid artefacts caused by osmotic shrinkage, samples were gradually dehydrated in glycerol (2-80%) and then ethanol (20 to 100%) and mounted in methyl salicylate (Sigma-Aldrich, M6752) between two No1 coverslips using custom spacers (stacked single [∼60 µm thick] and double-sided [∼100 µm] 3M Scotch Magic Tape) for imaging.

The low molecular weight of Neurobiotin allows it to cross gap junctions and cause a phenomenon of “dye-coupling”.^3,4^ We have not observed this phenomenon, suggesting that the neurons investigated in this study do not have electrically connected partners.

### Imaging and rendering

Serial optical sections were obtained on a Zeiss 710 confocal microscope at 2 µm with a Plan-Apochromat 10x/0.45 NA objective, at 1 µm with a LD-LCI 25x/0.8 NA objective, and at 1, 0.5 or 0.3 µm when a Plan-Apochromat 40x/0.8 NA objective was used. DyLight 488 NeutrAvidin and Texas Red phalloidin-treated samples were imaged using 488 and 594 nm lasers, respectively. Images were processed in Fiji (http://fiji.sc/) and Photoshop (Adobe Systems Inc.).

### Identification of wing and neck muscles MNs and TSDNs

The motoneurons were unambiguously identified via the choice of the back-fill’s point of entry. The identity of individual TSDNs was concluded from following anatomical features: position of the cell body and the morphology of the arborisations in the brain; morphology of the processes in T2, T3 and A1; and position/location of the interneuron’s main axon in the connective. We used work by Olberg^5^ and Gonzales-Bellido^6^ as a reference. Although the work of those authors was conducted on *Aeshna umbrosa* and *Anax junius* (Olberg) and *Libellula luctuosa* (Gonzales-Bellido), we found the level of morphological conservation to be high enough to identify individual TSDNs with good confidence. The nomenclature of muscles and their motoneurons was adapted from Clark.^7^ This neuroanatomical study is based on the analysis of more than 150 ganglionic/CNS preparations.

### Stimulus generation and projector setup

A modified DepthQ highspeed projector and StimGL software were used for producing the visual stimulus^6^. We operated the projector at 1280 x 720 pixels at 360Hz, using an external blue light source (SugarCUBE™ LED Illuminator) to drive the DLP. A light intensity mask was required to make the projected image on a flat screen more homogeneous. This mask was generated based on the measurement from a LMK Mobile Imaging Photometer (Opteema Engineering GmbH), and loaded into StimGL. A flat back-projection screen made of vellum tracing paper was placed 6cm from the head of the dragonfly. The projector setup was configured to run in two modes (Figure 1B). For the angular size-speed mapping, the projection covered ∼80° in elevation and ∼45° in azimuth covering the center and the dragonfly’s right receptive field (14.3 ∼ 24.5 pix/°, depending on eccentricity from the center). For the naturalistic target playback experiment, the projection covered ±60° in azimuth and ∼80° in elevation (6.1 ∼ 24.5 pix/°). These pixel densities guaranteed that even the smallest target still could be projected, given the limitations of the projector. To account for the reduced screen occupancy for highspeed targets, we normalized all size-speed mapping responses by the target screen occupancy.

For wide-field motion stimuli, a directional grating function with ∼5° spatial period was presented at the center of the visual field without perspective correction in accordance with previous experimental protocols.^5,8^ For experimental stimuli, on the other hand, the angular size and speed of the projected target had to be controlled on the flat screen. This included the appropriate aspect ratio and rotation correction of the circular target given the fixed head position of the dragonfly relative to the screen. A circular target (1° in size, 100°/s) moving across the center of the dragonfly’s visual field in 12 directions in pseudo-random order was used as a standard test stimulus to help detect candidate TSDN units when inserting the electrode.

### Electrophysiology for acute recordings

A temperature-controlled dragonfly holder was used following the procedure described in.^6^ This device allowed stable control of the dragonfly’s thoracic temperature and additonially provided a reliable head restraint. The thoracic temperature was verified with a thermocouple inserted inside the thoracic cavity in a few preparations with the holder temperature at a steady state. After the animal was immobilized and cooled to 8∼10°C, an incision was made between the pro-thoracic and meso-thoracic segment on the ventral side of a restrained dragonfly. An air sac was removed or pushed aside to allow access to the ventral nerve cord (VNC). A tungsten electrode (3Mohm, A-M Systems, Inc.) controlled by an MP-285 Micromanipulator (Sutter Instrument, Inc.) was inserted through the sheath of the VNC. While the standard test stimulus was playing from the projector, fine adjustments to the electrode position were made until a high amplitude and well isolated spike with a clear target response was attained. We then confirmed for the lack of response to wide-field motion using the moving grating stimulus in 8 directions. Finally, we checked the oscilloscope spike triggered overlay to make sure the signal-to-noise ratio (SNR) was at least 5 for unit isolation. After these checks, we proceeded with the experimental stimuli (playback experiment or size-speed tuning, see main text for details). All acute extracellular recordings were made by an AM Systems 3600 amplifier and digitized with a National Instruments USB-6210 at 40kHz.

We recreated the immobilized studies but corrected the key conditions in two experiments to separately probe the effects of temperature and prey motion on TSDN responses. In the first experiment (n=5 dragonflies), we systematically varied the dragonfly’s temperature from torpor to flight temperatures (16°C, 20°C, 26°C, 32°C). We used a custom projection system capable of producing moving stimuli that spanned the full range of prey angular sizes and speeds that have been measured behaviorally^2^ (Figure 1B). The visual stimulus consists of a constant angular size target traveling at constant angular velocity in two orthogonal directions. Instead of mapping the spatial receptive field, we focused on mapping the target angular state tuning. Across trials, target size and speed were varied randomly, to fully sample the target angular size-speed parameter space. Two different angular size ranges were explored: a broad range (1∼10° in 0.2° steps) as in the original TSDN experiments^5^, and a narrow range (0.1∼1° in 0.1° steps) that reproduces the angular prey sizes seen prior to takeoff in real interception flights^2^. Each mapping session consists of 20 repetitions of 200 combinations of angular size-speed. Given the temperature treatments, repetitions, two angular size ranges, and stimulus intermissions, each animal was subjected to a minimum of 4hr recording session. Thus, only 5 dragonflies successfully completed the size-speed mapping. We functionally isolated a vertically sensitive TSDN from 2 dragonflies, a horizontally sensitive TSDN from another 2 dragonflies, and both from the fifth dragonfly. To pool data across dragonflies, we normalized the spike counts and plotted the firing probabilities of TSDNs as a two-dimensional tuning function (Figure 2A-B, D-E). In general, TSDNs spike in bursts and often with 2∼3ms intervals (Figure 2C,F). The temperature impacted the number of spikes and the length of the spike trains (Figure S2). In the second experiment in which realistic head-centered stimuli were used, the TSDN’s inter-spike interval histogram is comparable to those from the size-speed mapping experiment with peaks at ∼2ms and ∼4ms (Figure 3F). These sparse, bursty responses are indeed a characteristic of TSDNs and their putative upstream partners STMDs.^9^

### Wireless TSDN recordings from a freely behaving dragonfly

#### Basic principles

The neural telemetry system is based on a custom chip, flex circuit, and RF power harvesting system described in.^10^ It contains 10 neural channels and 4 EMG channels with a bandpass filter (250Hz∼10kHz) and 200x gain. A custom made 915MHz dipole antenna was used to harvest 20W RF energy from a base station (Figure 1C) to power the amplifier circuit, ADC and an RFID communication circuit. The digitized signals were encoded through modulating the impedance of the dipole antenna, which can be detected by the base station as a backscattered signal. For details of the system please refer to the instrumentation paper.^10^

To use neural telemetry to record from neurons in the flying dragonfly engaged in prey capture required solving several key technical challenges (Figure S1A-B). First, the electrode implantation must not compromise the flight behavior both physically (e.g. joint mobility) and physiologically (e.g. neuronal tissue damage). Second, the weight of the backpack must be kept extremely low to minimize the impact on flight maneuvers necessary for interception flight. Third, the antenna must be crash-resistant as the freely flying dragonfly could potentially collide with the artificial target system or any experimental equipment inside the flight arena. Fourth, the telemetry backpack mounting must not interfere with the flapping wings or shift the centre of mass significantly. Fifth, it must be integrated with the motion capture system so the neural recordings can be time-locked to the behavior. Finally, the system must be carefully tuned to allow adequate range for good data transmission.

The bare die of the telemetry chip was wire bonded onto a flexible PCB with shielding epoxy. The full implanted telemetry backpack weighed ∼38 mg, including the antenna. The motion capture marker frame (to track the dragonfly’s body orientation), and glue brought the total weight to ∼45 mg (Figure S1B). A custom bi-polar electrode was fabricated by insulating 0.0005” bare gold wires (California Fine Wires Co.) with polyimide tubing also acting as the shaft. The length and orientation of the exposed gold wires were trimmed manually to obtain an impedance between 100kΩ ∼ 500kΩ. The recording electrode and backpack were implanted separately and then connected to each other, during a procedure described below. The night for implantation, dragonflies in torpor were removed from the flight arena (∼8pm) and iced for 15 minutes before being inserted into a custom mounting apparatus that immobilized the dragonfly and oriented it for surgery (Figure S1B). The implantation and backpack mounting process took 3∼4hr. If the implantation was successful, the dragonfly was placed back into the flight arena the same evening, and behavioral experiments began the following day. As soon as the electrode was implanted, the animal’s health degraded gradually. However, the dragonfly would not hunt immediately after surgery. Operating on the dragonfly the evening prior to experiment was found to be the optimal compromise.

The dragonfly was monitored at the first light of artificial dawn in the dragonfly arena at 8am the next morning. The artificial prey system, telemetry base station, and motion capture system were checked and prepared to commence experiments no later than 10am, when the flight arena temperature reached a steady state (32°C). By this time, the dragonfly would have taken some morning flights and started to find a perch for breakfast. Typically, the instrumented dragonfly behaved normally for the first day but started to weaken during the second day. As a result, all experiments only ran for one day.

#### Electrode implantation and backpack installation process

We used a single bipolar electrode wired to 2 channels for a duplex sampling rate of 56KHz. The electrode was implanted near the nerve cord through a ∼250um hole inserted into a soft tissue joint between thoracic segments T1 and T2, on the ventral side of the animal’s body slightly below the front legs (Figure S1A). This recording site ensured that the largest descending neurons near the electrode were TSDNs and would be the primary signal measured. The anatomical location itself prevented recording from motoneurons to the neck or first pair of legs (which lie upstream) or wings (which lie downstream in the thoracic ganglion).

The procedure for obtaining TSDN responses was the same as that described for acute recordings except that the incision and tissue removal had to be significantly minimized. In addition, a special dragonfly holder (Figure S1B) was used to align the telemetry backpack and to facilitate wiring on the dragonfly’s body. The holder did not support active cooling so dragonflies were pre-chilled on ice before the surgery began. Once a clear directional response was obtained, the shank of the electrode was glued in place around the cuticle incision (LOCTITE 4305 UV adhesive, Henkel Ltda). This process required iteratively building and curing traces of glue with a UV gun controlled light spot promptly. After the electrode was glued to the exoskeleton, micro scissors were used to cut off the supporting glass manipulator and the wire tether, releasing the electrode from the rack-mounted amplifier. At this point, the preparation was transferred to a microscope-based assembling station to solder the electrode to the telemetry backpack (Figure S1B).

The 3D-printed dragonfly holder could position the pre-tested telemetry backpack to the back of the dragonfly’s thorax to ensure proper alignment to the anatomical structures (Figure S1B). A thin layer of superglue was used to prepare the surface of the cuticle. Then the backpack was secured to the thorax using UV glue. Soldering the electrodes wires directly was unwise, as these thin gold wires evaporated easily upon contact with the soldering iron. Instead, the gold wires were wrapped around a thicker silver wire (0.003”, A-M Systems) first before a thin coat of solder was applied quickly to make the connection. The silver wires from the electrode were carefully bent and routed to the telemetry backpack, tacked down with small dots of UV glue. Soldering the silver wires to the backpack was another high-risk procedure as the heat from the soldering iron could easily damage the backpack electronics, or the dragonfly itself. However, we found the mechanical resilience of solder much more reliable than conductive adhesives like silver epoxy. Given good micro-soldering experience, the connection was feasible. The instrumented dragonfly typically gained ∼50mg in weight (including backpack, antenna, motion capture frame, and adhesives).

Why did telemetry dragonflies have low takeoff rates during predation? Inspection of the mean foveation error across head saccade events (n=55) shows that in many cases experimental animals struggled to maintain prey foveation in the critical pre-takeoff period. While head saccades of telemetry dragonflies had comparable dynamics of normal animals that pursued prey, foveation error was consistent with normal dragonflies that started the pre-takeoff process but ultimately did not take flight (Figure S3C). In many cases, the experimental animals were not able to maintain or gave up foveation at 100ms post-saccade. This suggests that the resting position of the head and the general target detection mechanism were not affected by the neural implant. Instead, neural tissue damage, the presence of the implanted electrode, and the presence of head mounted motion capture markers collectively contributed to the lower takeoff rate. As a result, we collected neural telemetry data for pre-takeoff behavior from dragonflies with head markers (n=3), and data for prey interception flights from dragonflies without head markers (n=3).

### Spike sorting

For acute extracellular recordings, we aimed to isolate the units of interest with electrodes with impedance from 2Mohm to 3Mohm. Thus, the resulting signal typically contained up to 3 units with one unit clearly biggest even when the body temperature was raised to 32°C. These recordings did not require spike sorting as we could simply use an amplitude threshold to extract the large unit we isolated. For neural telemetry recordings, the signal was more complex due to lower impedance (∼150kΩ), motion artefacts not cancelled by the differential channel, and significantly more background neural activity during behavior. Since TSDNs fired sparsely, and the total available recordings were limited, automatic spike sorting algorithms such as KiloSort2^11^ did not perform well and lost many valid spikes. To sort units, we thus applied the most conservative and deterministic approach: template matching. Candidate spikes were picked out manually to form templates which were used to search through the same recording session. A template deviation index was computed for every spike peak-aligned to the template by integrating the deviation over ±0.5ms around the negative peak (Figure S4A). To allow comparison across different templates and to account for the diminishing importance away from the peak, we normalized deviation (relative to the standard deviation of the template) and applied a custom weighing function before the integration. In short, a low template deviation index means the spike in question matches the template well especially around the peak. We used this index to help with manual spike-sorting in the neural telemetry dataset. Each putative unit was checked for contamination though the inter-spike-interval (ISI) histogram (Figure S4B-C). TSDNs have very short refractory period and sometime produce spike duplets under 1ms interval at >30°C (Figure S1C). Thus, we isolated spikes with ISI below 1ms from any unit first and manually reallocate them back to the unit as appropriate. The entire template-matching process was later integrated into a MATLAB Graphical User Interface (GUI) as an App (https://github.com/dragonflyneuro/NBits-Dragonsort; app developed by Dr Daniel Ko) for better user experience. However, as noted in the main body of the paper, we are not attempting to assign these signals to identified TSDNs, and whether the signals are pure single units or multi-unit mixes does not affect the results.

### Neural telemetry data processing

We identified TSDNs in behaving dragonflies through several mechanisms. The small target motion sensitivity and directional receptive fields of putative cells were assessed during the surgical procedure (see Methods). This ensured the electrode was implanted in the correct location, adjacent to TSDN axons in the nerve cord. All implantation was performed in the evening to allow the dragonfly to recover over one night’s sleep. The telemetry experiments were conducted on the day after implantation (within 8∼16hr post implantation). The neural waveforms often drifted overnight, and new units appeared when the animal woke up. Unfortunately, there was no practical way to perform systematic receptive field mapping once the dragonfly was freely flying. Recapturing the dragonfly or any intervention would only irritate the insect and risked the loss of normal behavior (i.e. going into escape mode). Thus, we could not use receptive fields to identify TSDNs.

Given the characteristics of TSDNs and how we performed the implant, target response was defined by three criteria: **1)** presence without head movements; **2)** sufficient amplitude and compatible inter-spike-interval; **3)** exhibit direction selectivity to sub-degree moving targets. These criteria ruled out motion artifacts, generic motion sensitive neurons, anatomically irrelevant units, and strong multi-unit signals. In practice, we concatenated 25∼30 trials evenly across the recording session that did not exhibit any motor activities (i.e. head or body) and contained an equal representation of inbound and outbound artificial target runs. These trials served as the seed dataset (type 1 data; Figure S4B) to form spike templates for each recording session, satisfying criteria #1. From this dataset, only spikes with signal-to-noise ratio of at least 3 were considered (criteria #2). Finally, we excluded spiking instances when the artificial target was outside the prey detection saccade cone with 40ms time margin. The above process generated a handful of spike templates (Figure S4B), which we then use to search for spikes in all other trials with head movements (type 2 data; Figure S4C). For units that exist in both types of trials, we computed the direction selectivity by comparing the spike counts corresponding to artificial targets going in opposite directions (Figure S4B-C, criteria #3). Given the target direction presented, we then defined units with over 60% spikes in a 60° directional polar interval as directional selective. For dragonflies without head markers, the first criterion was modified to be: presence in both non-takeoff and takeoff trials. We did not sort TSDNs into identified units (e.g. MDT1), and none of the results to follow are dependent on the cell-type specific TSDN nor on whether the unit was perfectly isolated or partially multi-unit.

### Kinematics data collection and processing

The neural telemetry backpack installed on the dragonfly incorporates a carbon marker frame for three-dimensional motion capture (Figure S1A). For some dragonflies, we also installed head markers for reconstructing the head orientation during behavior (Figure S1A). The 3D kinematics procedures follow our previous work.^1,2^ We used a combination of real prey and artificial prey. The artificial prey delivery system consisted of a motorized pulley system dragging a small bead supported by two supporting poles.^2^ We could assign the system to drive a variety of constant speed targets at specific heights. Due to the low rate of takeoff from instrumented dragonflies, we restricted the equivalent target angular speed to under 500°/s and angular size to under 0.7° to facilitate target tracking.

### Size-speed map production

To systematically map the size-speed tuning of TSDNs. We made a perspective corrected target with fixed angular size and angular speed moving either in the fore-aft vertical direction or left-right lateral direction. To reduce habituation, an intermission of 3∼5s was introduced between angular size-speed state presentation. Furthermore, the presentation sequence was randomized, and the target trajectories were offset by ±5° randomly. Due to the high density of state space sampling, each state was played only once as similar states would be sampled nearby. To pool spike count data across animals, we divided each size-speed state bin by the total spike count for the corresponding mapping recording. This normalized activity map was then averaged across animals and converted to % activation. To control for slow targets remaining on the screen longer, we finally divide the size-speed map by a weighing matrix derived from time in visual field.

For size-speed mapping of freely behaving dragonflies, we compiled all target presentation samples within the perched dragonfly’s saccade cone to construct a stimulus prior map for each animal. Then we counted spikes that occurred when the target was within the same saccade cone. For trials with an foveating head saccade, we counted spike prior to the head movement. Each spike was associated with the target size-speed 30ms before the spike time, based on the published TSDN response latency. The aggregate of these spike states formed the preliminary size-speed map. Since we have fewer spikes than the restrained recordings, we performed a weighted average based on the total spike count to merge data across animals. The average size-speed map was divided by the aggregate prior map to obtain the activation rate as the number of spikes per second of stimulus presentation.

To recreate natural prey trajectories we computed the angular size and speed from a dataset of dragonfly (*Plathemis lydia*) intercepting fruit flies (*Drosophila virulus*)^1^ in two versions. The first was computed with a coordinate system referenced to the dragonfly’s perch location (perch-centered), simulating the prey statistics prior to takeoff (Figure 2G-H, cyan). The second version was computed from a moving reference frame reflecting the dragonflies time-varying inflight location relative to the prey, as seen in prey interception flights (Figure 2G-H, blue). To examine the effect of head movement, we computed the prey size-speed state in both the body and the head coordinate system (Figure S3A-B) from another dataset of dragonfly (*Plathemis lydia*) intercepting artificial prey.^2^ The foveating head saccades caused a large angular speed fluctuation prior to takeoff (Figure S3A-B). This pushed the target state temporarily outside the tuning range of TSDNs.

### Timing analyses

To define behaviorally relevant stages, we adopted the functional fovea, target detection saccade cone and the takeoff (tracking) cone from our previous study for the same species.^2^ To understand the timing of target responses within the target interception behavior, we computed spike time distribution across events relative to the onset of head saccade, the time of takeoff, and the time of zenith crossing. The zenith crossing was defined as the time when the target arrived at its closest proximity to the zenith line directly above the dragonfly’s perch location. For the playback experiments, the virtual saccade and takeoff corresponded to the time of these events in the behavior data from which the stimuli were derived.

---

**Figure S1.**
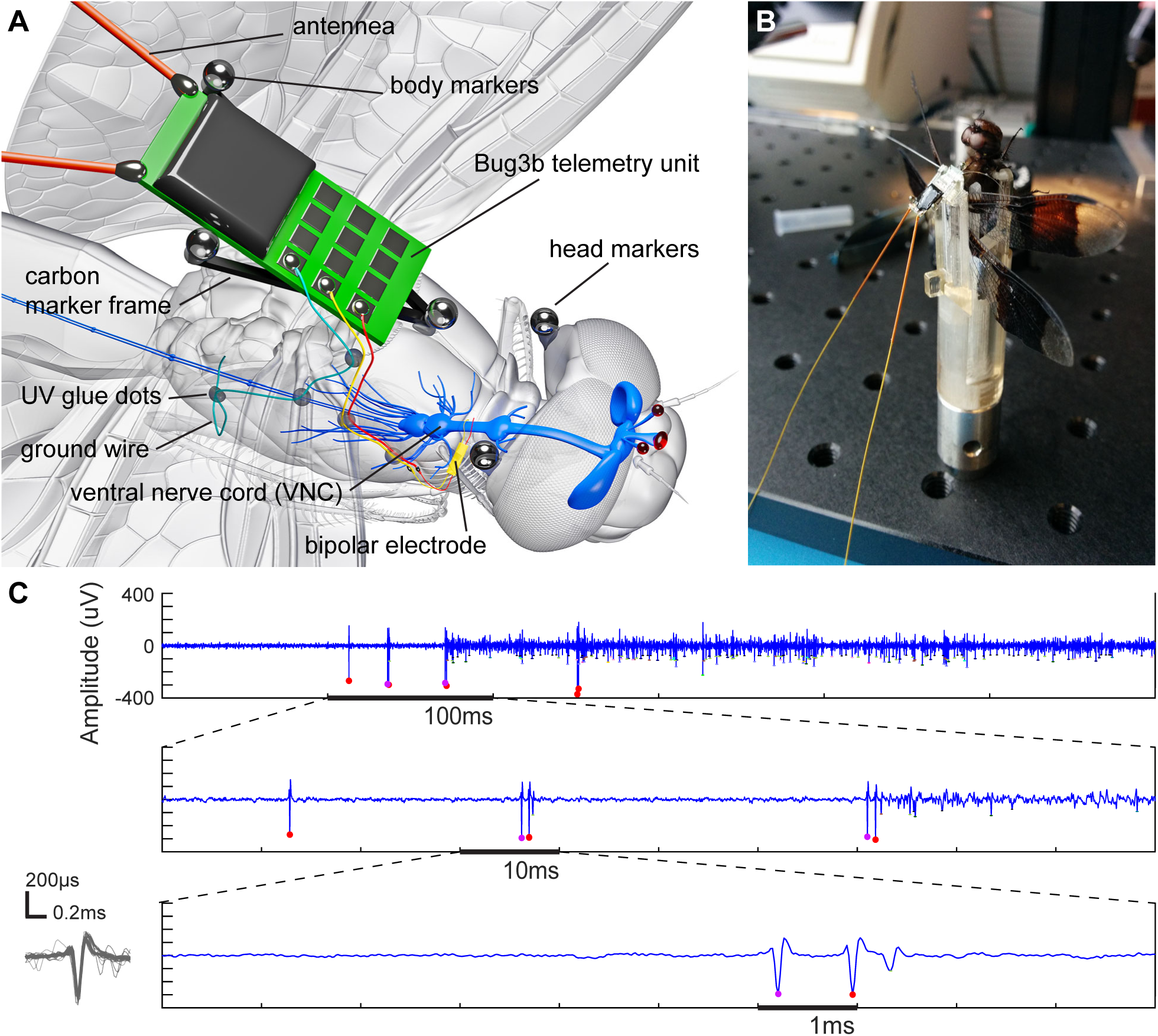
Dragonfly neural telemetry implantation. **A)** The full neural telemetry installation starts with a custom bipolar electrode latching onto the neck connective between T1 and T2 of VNC. The insulating polyimid tubing extended just outside of the body and was secured on the cuticle with sealing glue. The gold electrode wires transitioned into larger silver wires for connection to the telemetry backpack. The teletemetry backpack was secured precisely on the carbon marker frame for motion capture. **B)** The telemetry backpack measures 9.5mm x 3.9mm x 0.8mm and weighs ∼38mg. The flexible antenna maintains a geometry that does not interfere with wing motion but still allows acceptable RF performance. A custom dragonfly restraining holder was designed and fabricated in order to allow the multi-stage process of electrode implantation and telemetry backpack installation. **C)** A successful backpack installation produced good quality data (56kHz sampling rate) from actively hunting dragonflies. This example came from a perched dragonfly watching a potential prey. The TSDNs exhibit high firing dynamic with spike duplets (marked with magenta/purple dots for visualisation in this example). After a handfull of TSDN spikes, some motor-like background signals are correlated to the active target tracking head movements. The characteristic duplexes sometimes have an interval below 1ms, consistent with the inter-spike interval distribution in Figure 2**-3**.

**Figure S2:**
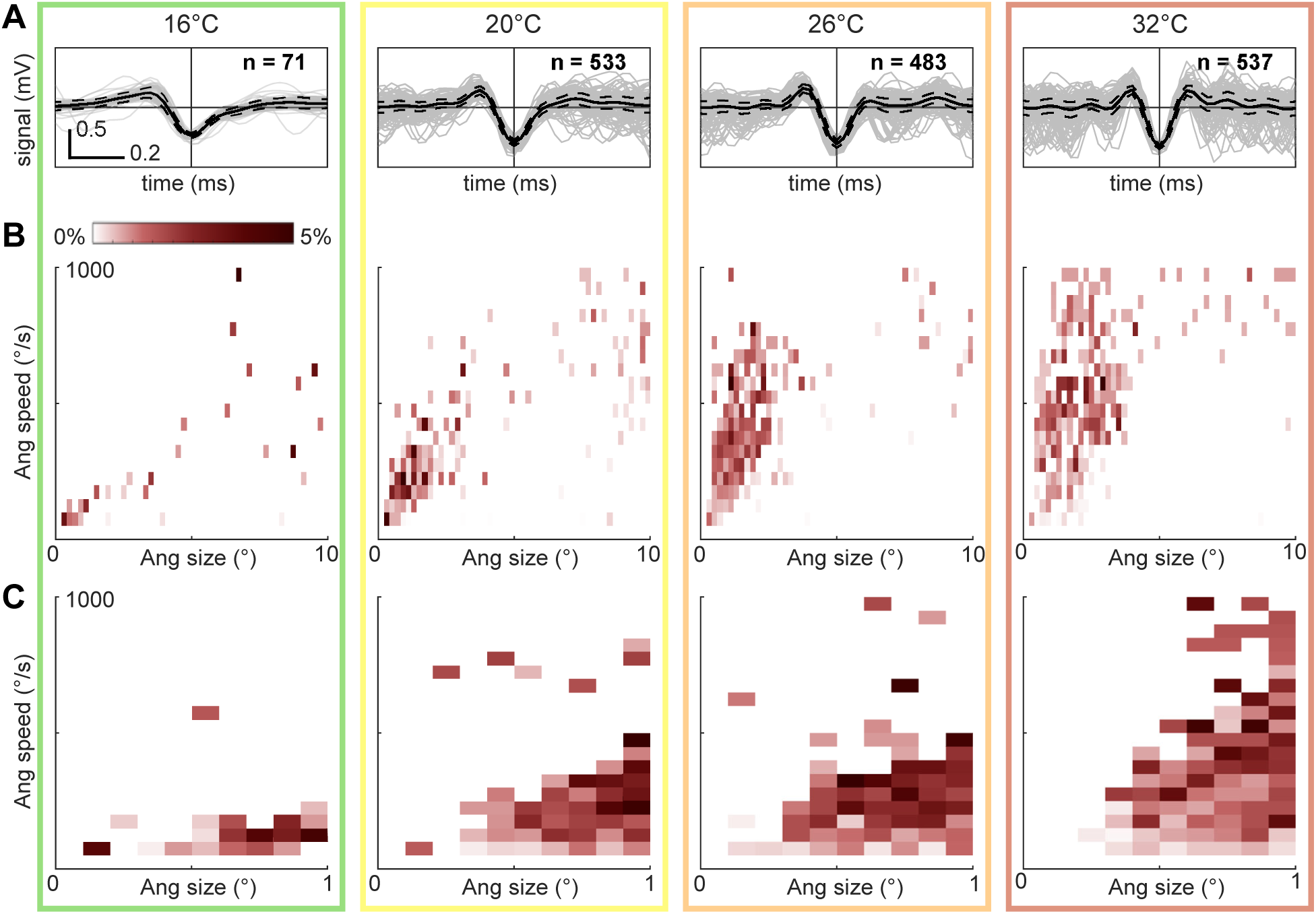
The effect of body temperature on TSDN target size-speed tuning. **A)** The same unit spike overlay across four different thoracic temperatures (color matched to Figure 2C**,F**), with mean (black solid line) and standard deviation (dashed lines) marked. Given increasing temperature, TSDN spike width shortened significantly from ∼0.5ms at 16°C to ∼0.25ms at 32°C. **B)** The size-speed response mapping for the same unit across temperatures at the 0∼10° angular size range. To account for very different spike counts, the tuning maps were shown as normalized firing % activation. **C)** The same unit was independently re-mapped with target size below 1°. With increasing temperature, the cell responded to higher angular speed targets. 32° is the most naturalistic body temperature for hunting *Plathemis lydia*, and is the most consistent with prey selection and telemetry recordings (Figure 2I).

**Figure S3.**
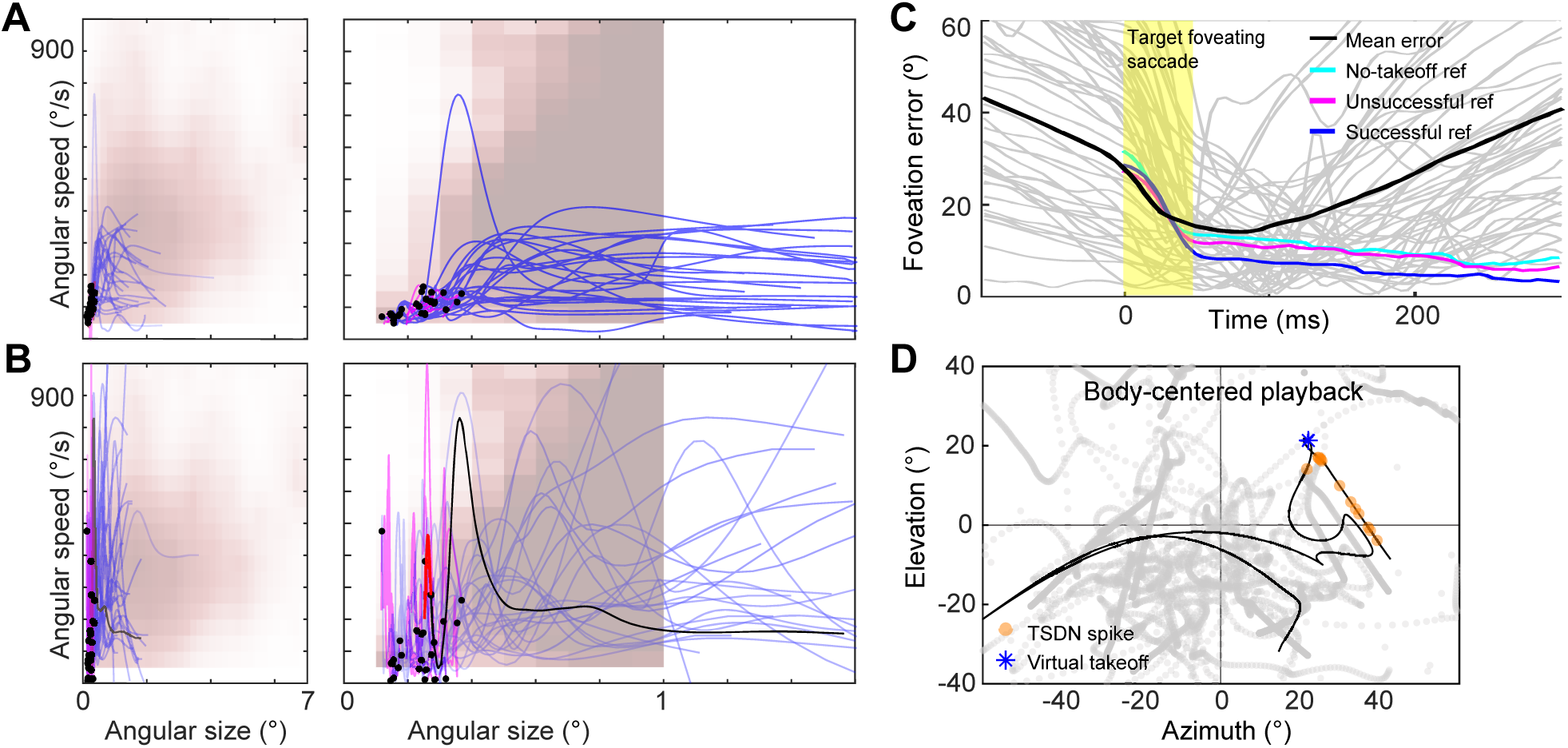
Target visual state evaluation, foveation perofrmance, and body-centered playback stimuli. **A)** Artificial target size-speed state evolution during real interception flight stays well-within the TSDN tuning with respect to the body reference frame. Pre-takeoff segments are in magenta. Blakc dots mark the moments of takeoff. **B)** The same data in the head reference frame shows that the initial 50ms head saccade motion pushed the angular speed outside the TSDN tuning temporarily. Afterward, the angular speed stays mostly within the TSDN tuning. **C)** Dragonflies with telemetry backpack and head markers (3 dragonflies, n=155 events, grey lines) performed foveation. The mean foveation error (black line) rapidly drops below 20° with the foveating head saccade, but quickly rises again in 50ms as the aniaml gives up or fails to maintain tracking. To provide a comparison, the performance references are reproduced from our previous study (**Lin & Leonardo 2017**) in which we categorized target interception trials as successful flight, unsuccessful flight, no-takeoff head tracking. In the previous study, the dragonflies only had body and head markers mounted. **D)** Same data in Figure 3B projected to the body-centered reference frame. Target size variation was not shown for visualization purposes as we cannot see sub-degree targets. These stimuli emulate if the head was not actively tracking the prey, providing a baseline contrast to the effect of head foveation. This type of stimuli is analogous to the ones used in **Ogawa *et al* 2023**. However, unlike the hoverfly study, the dragonfly performs more drastic maneuvres during interception flights. An example trajectory shows that after the virtual takeoff (blue *), the target moves with larger excursions.

**Figure S4.**
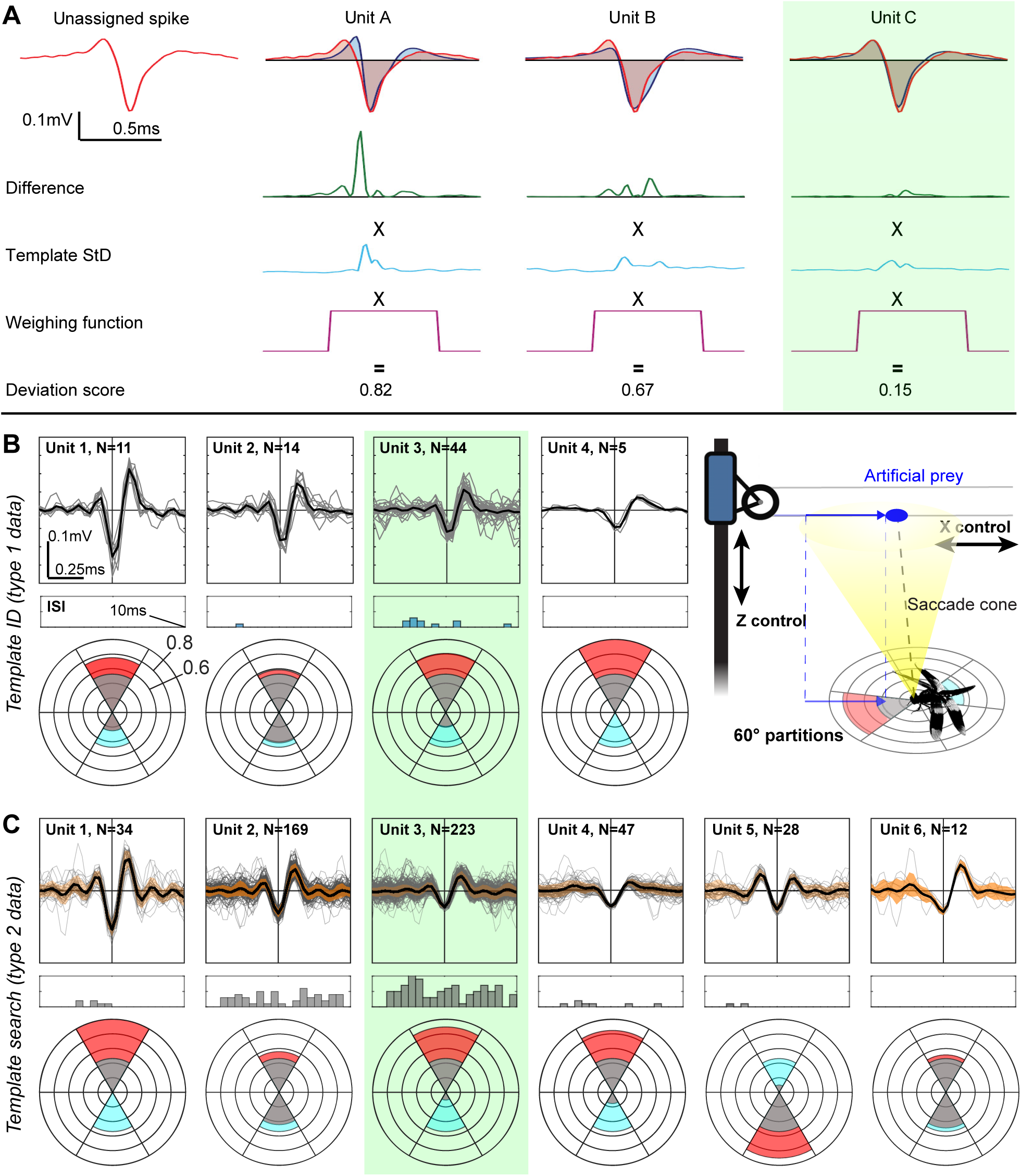
Spike sorting principle and target response identification from neural telemetry data. **A)** Any unassigned spike was compared to templates derived from the average waveform of manually identified units within a segment of the recording. The waveform difference was normalized and weighed to compute a deviation score. The spike was assigned to the unit with the lowest deviation score (green shaded unit). Illustration courtesy of Daniel Ko. **B)** Four units were identified from events without motor activities (type 1 data). The ISI was checked to minimize multi-unit condition (especially below 1ms range). The direction selectivity was computed for each unit (red polar) with the stimulus directional distribution as reference (cyan polar). Three of the four units were directional in this case. **C)** The same 4 units from B were used as templates to sort events with head foveating movements (type 2 data) from the same experiment. Standard deviation is shown in orange shade around the mean (black lines). The ISI distribution shows nothing under 1ms range in this example. The direction selectivity was consistent. Additional two new units were discovered. For spike timing analysis, we selected unit(s) that existed in both conditions, showed direction selectivity, and accumulated sufficient number of spikes (green shaded unit).

